# Preeclampsia in mice carrying fetuses with APOL1 risk variants

**DOI:** 10.1101/2024.03.20.586039

**Authors:** Teruhiko Yoshida, Khun Zaw Latt, Shashi Shrivastav, Huiyan Lu, Kimberly J. Reidy, Jennifer R. Charlton, Yongmei Zhao, Cheryl A. Winkler, Sandra E. Reznik, Avi Z. Rosenberg, Jeffrey B. Kopp

## Abstract

African-American women have a maternal mortality rate approximately three times higher than European-American women. This is partially due to hypertensive disorders of pregnancy, including preeclampsia. Fetal *APOL1* high-risk genotype increases preeclampsia risk, although mechanisms remain elusive. We characterized two mouse models to investigate whether fetal-origin APOL1 induces preeclampsia and which cell types contribute. We *in vitro* fertilized mice with sperm from two transgenic mouse lines: *APOL1* transgenic mice carrying human genomic locus constructs from bacterial artificial chromosomes (BAC) containing the *APOL1* gene, mimicking expression and function of human APOL1 (BAC/APOL1 mice) and albumin promoter *APOL1* transgenic mice expressing *APOL1* in liver and plasma (Alb/APOL1 mice). Dams carrying either BAC/APOL1-G1 or Alb/APOL1-G1 fetuses had elevated systolic blood pressure, while dams carrying BAC/APOL1-G0 or Alb/APOL1-G0 fetuses did not. BAC/APOL1-G1 and Alb/APOL1-G1 fetuses weighed less than littermates, indicating intrauterine growth restriction. Single-nucleus RNA-seq of APOL1-G1 placentas showed increased expression of osteopontin/Spp1, most prominently in vascular endothelial cells with robust APOL1 expression. Cell-cell interaction analysis indicated pro-inflammatory signaling between placental cells and maternal monocytes. These models show that fetal origin APOL1-G1 causes preeclampsia, inducing pro-inflammatory response in placenta and maternal monocytes. The APOL1-G1 variant poses a multi-generational problem, causing effects in mothers and offspring.

## Introduction

Preeclampsia is a leading cause of maternal and fetal mortality and morbidity, responsible for more than 76,000 maternal and 500,000 infant deaths per year worldwide.^1,2^ Preeclampsia is a multisystem disorder characterized by the new onset of hypertension, proteinuria, dysfunction in various organs, occurring during gestation or postpartum.^3^ The pathogenesis is thought to involve both abnormal placentation and maternal systemic vascular dysfunction, particularly involving endothelial dysfunction, although the underlying pathogenesis still remains elusive.^4,5^ Women with sub-Saharan ancestry have higher risk for hypertensive disorders of pregnancy including preeclampsia, compared with women with European ancestry.^6^ Despite advances in maternal care, African-American women continue to have a maternal mortality rate 2.6 times higher than European American women.^7^ Further, African-American newborns also have a mortality rate 2.4 times higher than European-American newborns (10.4 vs 4.4 deaths per 1,000 births) in their first year.^8^

*APOL1* high-risk genotypes, carriage of two *APOL1* high-risk variants (G1 or G2), pose a major genetic risk for kidney diseases in individuals with sub-Saharan African ancestry.^9,10^ Approximately six million individuals in the USA carry *APOL1* high-risk genotypes.^11^ Recently, the association between fetal carriage of *APOL1* high-risk genotype and preeclampsia is reported.^12–14^ Fetal *APOL1* genotype is also reported to be associated with altered fetal growth in term infants and preeclampsia in preterm infants.^15^ APOL1-G0 (low-risk variant) or APOL1-G2 (high-risk variant) transgenic mice under the control of nephrin gene (*Nphs1*) promotor develop preeclampsia.^16^ However, it is still not known if fetal *APOL1* high-risk variant causes preeclampsia.

We established and characterized mouse models of APOL1-associated preeclampsia, in order to investigate the role of fetal APOL1 variants during pregnancy. We used *in vitro* fertilization with sperm from two transgenic mouse lines, mimicking expression and function of human APOL1. The first set of transgenic mice (BAC/APOL1 mice) contained human genomic constructs from bacterial artificial chromosomes (BAC) containing APOL1-G0 or APOL1-G1. The second set of transgenic mice (Alb/APOL1 mice) had the mouse albumin promotor regulated expression of cDNAs encoding human APOL-G0 or APOL1-G1. We characterized gene expression of mouse placenta using single-nuclear transcriptomics. We report potential mechanisms of fetal APOL1 high-risk variant-induced preeclampsia.

## Results

### Pregnant mice carrying fetuses with APOL1-G1 showed preeclamptic phenotype

We have generated and characterized pregnant female mice by *in vitro* fertilization of sperm from BAC/APOL1-G0, BAC/APOL1-G1, Alb/APOL1-G0, and Alb/APOL1-G1 mice (**Figure 1A**). BAC/APOL1 mice have human APOL1 expression^17^ and Alb/APOL1 mice have APOL1 expression in plasma and liver.^18^ Dams carrying offspring with BAC/APOL1-G1 and Alb/APOL1-G1 had higher systolic blood pressure (158.4 mmHg [141.3-163.4], P=0.005 and 159.0 [137.3-164.8], P=0.0002), compared to dams carrying BAC/APOL1-G0 and Alb/APOL1-G0 mice did not (115.9 [114.5-130.8] and 109.5 [96.3-133.3]) (**Figure 1B**).

**Figure 1.**
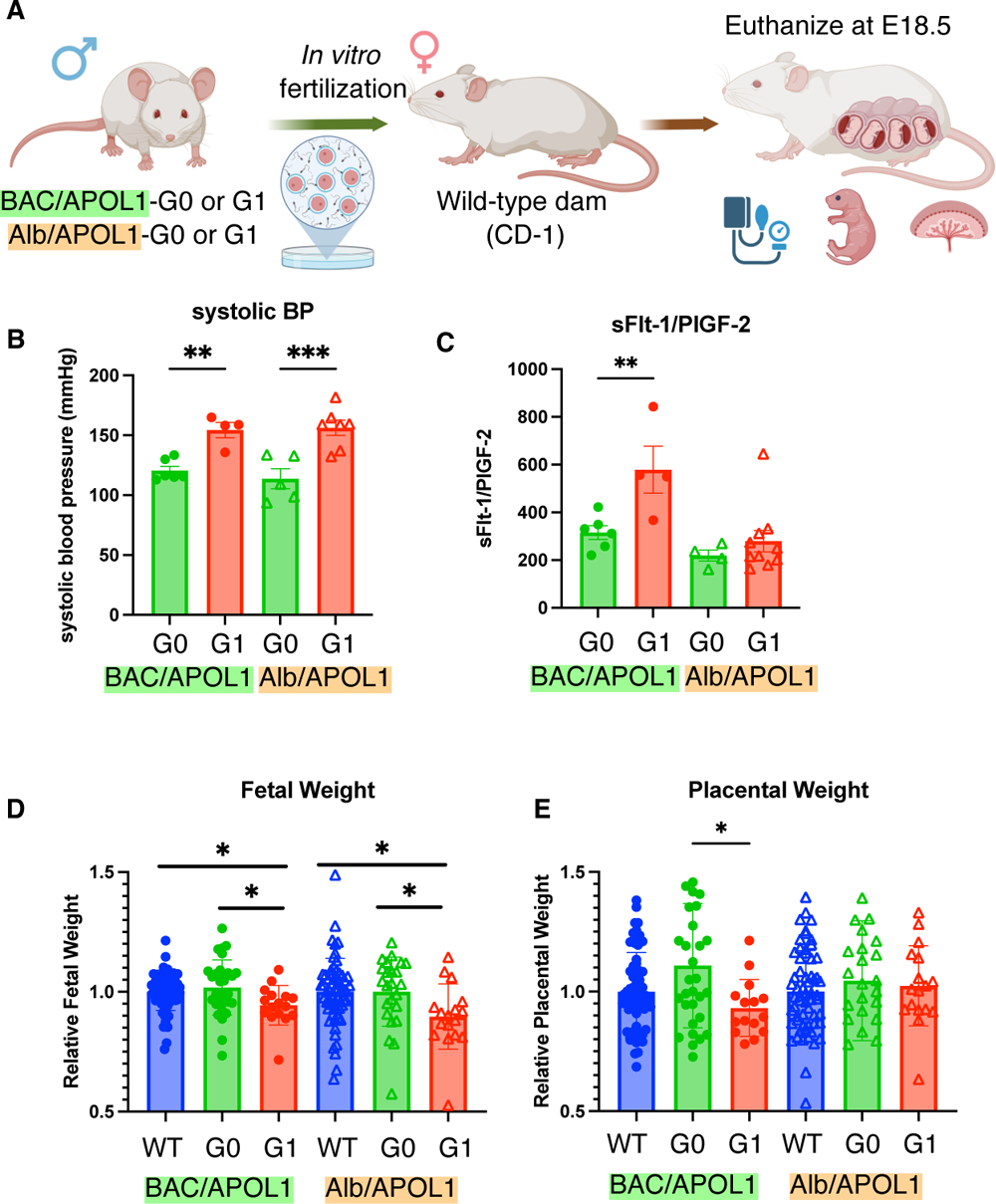
Preeclamptic phenotype was shown in dams carrying APOL1-G1 fetuses. (**A**) Overview of experiments. (**B**) Systolic blood pressure was elevated in dams carrying BAC/APOL1-G1 and Alb/APOL1-G1 fetuses, compared to dams carrying APOL1-G0 fetuses. (**C**) The ratio of sFlt-1 and PlGF-2 was elevated in dams carrying BAC/APOL1-G1 fetuses. (**D**) Fetuses with BAC/APOL1-G1 and Alb/APOL1-G1 were smaller in weight at E18.5, indicating intrauterine growth restriction. (E) Placenta with BAC/APOL1-G1 fetuses were smaller in weight.

With regards to preeclampsia markers, dams carrying BAC/APOL1-G1 fetuses had higher sFlt-1/PlGF-2 ratio compared with dams carrying BAC/APOL1-G0, although we did not see the same trend in dams carrying Alb/APOL1 fetuses (**Figure 1C**). Fetuses with BAC/APOL1-G1 and Alb/APOL1-G1 genotypes had smaller body weight (relative weight 0.95 [0.90-0.98], P= 0.02 and 0.90 [0.82-0.95], P=0.02) compared with fetuses with wild-type genotype, although we did not see the difference in fetuses with BAC/APOL1-G0 and Alb/APOL1-G0 genotypes compared with fetuses with wild-type (**Figure 1D**). Placentas with the BAC/APOL1-G1 genotype had a smaller weight compared with placentas with the BAC/APOL1-G0 genotype (**Figure 1E**). We did not see differences in serum creatinine and in albumin/creatinine ratio between genotypes (**Figure S1A, S1B**). Blood urea nitrogen, liver transaminases, and pathologies of the placentas and maternal kidneys did not show differences between genotypes (**Figure S1C-H**). Both BAC/APOL1 and Alb/APOL1 models showed that fetal APOL1-G1 induced preeclampsia with gestational hypertension and intrauterine growth restriction, indicating placental dysfunction.

### Single-nucleus RNA-seq from mouse placentas showed distinct cell types

To elucidate the molecular mechanisms dysregulated in placenta by fetal APOL1 genotype, we conducted single-nucleus RNA-seq of placenta as shown (**Figure 2A**). After pre-processing, 46473 nuclei remained and showed 24 distinct clusters by unbiased clustering (**Figure S2A**). We limited the data for the analysis to 37083 nuclei within 16 distinct clusters by removing cell clusters of fetal mesenchyme, doublet clusters and unknown cell populations as shown (**Figure 2B**). We annotated cell types by canonical markers based on a paper with the similar single-nucleus RNA-seq data of mouse placenta^19^ (**Figure 2C**). We confirmed that each sample contributed to each cell types, and the ratios of cell types were shown (**Figure S3A**). We conducted snRNA-seq using placentas of male offspring to identify cell origin by expression of a sex-dependent gene, *Xist*. Decidual cells and monocytes showed robust *Xist* expression, indicating these cells were mainly of maternal origin (**Figure 2C**). Multiple cell types in BAC/APOL1-G0 or G1 placenta expressed APOL1, including spongiotrophoblast (SpT), syncytiotrophoblast-2 (SynT2), glycogen cells, smooth muscle cells and Hoffbauer cells, with the highest expression in vascular endothelia as reported in other organs like kidneys (**Figure 2C, S3B**). There was low expression of *APOL1* in Alb/APOL1-G0 and -G1 placentas (**Figure S3B**).

**Figure 2.**
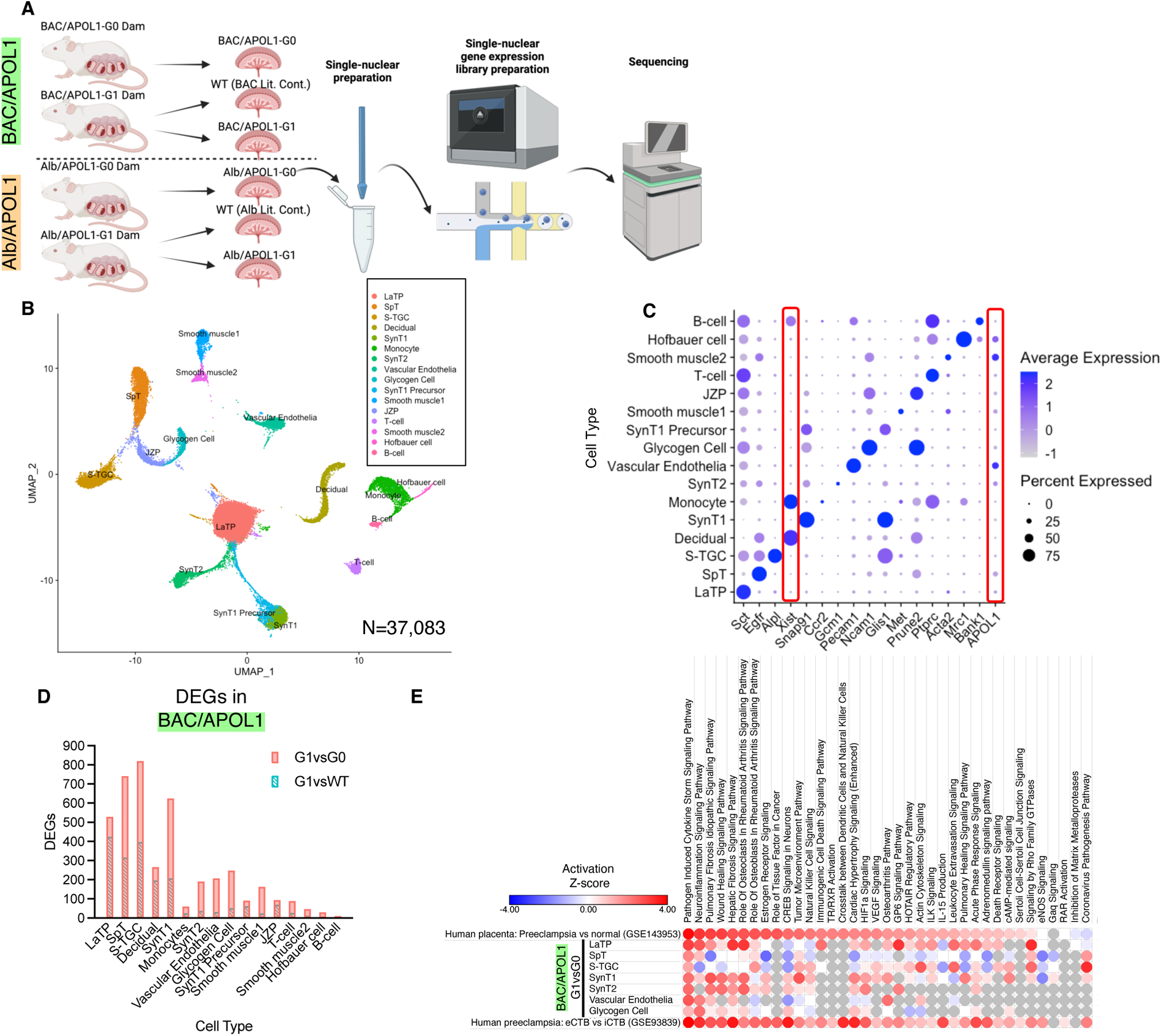
Single-nuclear RNA-seq of placenta profiled differential expressed genes in each cell types between APOL1 genotypes. (**A**) Overview of single-nuclear RNA-seq experiments (**B**) UMAP plot of single-nuclear RNA-seq data for analysis from six samples with 37,083 cells, showing 16 clusters. (**C**) Dot plot of marker genes characteristic for each cluster (**D**) Number of differentially expressed genes (DEGs) in each cluster, comparing APOL1 genotype (G1 vs G0 and G1 vs WT) in BAC/APOL1 model. (**E**) Heatmap showing canonical pathway activity z-scores of each cell type comparing BAC/APOL1-G1 and G0, human placenta comparing preeclampsia case from healthy case (GSE143953), and micro dissected human placenta comparing endovascular cytotrophoblast (eCTB) and interstitial cytotrophoblast (iCTB) (GSE93839). LaTP: labyrinth trophoblast progenitor; SpT: spongiotrophoblasts; S-TGC: sinusoidal trophoblast giant cells; SynT1: outermost layer syncytiotrophoblasts SynT2: intermediate layer syncytiotrophoblasts; JZP; junctional zone precursors

### Differentially expressed gene analysis showed inflammatory and human preeclampsia signatures in APOL1-G1 placenta

Differentially expressed genes (DEGs) were identified in each cell type comparing between APOL1-G1 and APOL1-G0 or APOL1-G1 and wild-type (G1 littermate control) in each mouse model (**Figure 2D, S3C**). As expected, a comparison between G1 vs G0 showed more DEGs than in G1 vs WT in the BAC/APOL1 model. Further, we compared DEGs of G1 vs G0 in each model. When 802 DEGs in SpT from BAC/APOL1 placenta were compared with 291 DEGs from Alb/APOL1 placenta, 116 DEGs were overlapped (**Figure S3D**). DEGs in vascular endothelia and monocytes were also compared and showing shared DEGs in both models (**Figure S3E and S3F**).

Canonical pathways activated in each cell type comparing BAC/APOL1-G1 and G0 were analyzed. Activated pathways in BAC/APOL1-G1 placenta include inflammatory and autoimmune disease related pathways such as pathogen induced cytokine storm pathway, pulmonary fibrosis idiopathic signaling pathway, and HIF1α pathway (**Figure 2E**). HIF1α^20^ is involved in preeclampsia pathogenesis. To compare our model with human relevant data, we compared our data with published human placenta RNA-seq data comparing preeclampsia case and normal control (GSE143953)^21^, and we found similarities in activated pathways as shown (**Figure 2E**). This indicated that cells in BAC/APOL1-G1 placenta recapitulated human placental transcriptome with preeclampsia. Furthermore, we compared our BAC/APOL1 mouse data with microarray data from micro dissected placenta with preeclampsia to compare interstitial cytotrophoblast (iCTB) and endovascular cytotrophoblast (eCTB) (GSE93839).^22^ The comparison results suggested that pathways activated in LaTP, which is equivalent to human cytotrophoblast, from BAC/APOL1-G1 compared with G0 were more similar to eCTB compared with iCTB. Since iCTB is more invasive than eCTB, this indicated less invasiveness of LaTP in BAC/APOL1-G1 placenta, which may suggest that poor labyrinth formation could lead to this preeclamptic phenotype.

### Cell-cell interaction analysis showed activation of osteopontin/Spp1 pathway in APOL1-G1 placenta

We conducted cell-cell interaction analysis in each sample to identify differences in ligand-receptor interactions associated with the APOL1 genotype (**Figure 3A**). Overall, APOL1-G1 placenta showed more interactions compared with APOL1-G0 placenta in the BAC/APOL1 model (**Figure S4A and S4B**). When we plot all potential communication strengths in a 2D plot, multiple cell types in BAC/APOL1-G1, especially monocytes expressed more outgoing signals with ligands (**Figure S4C, S4D**).

**Figure 3.**
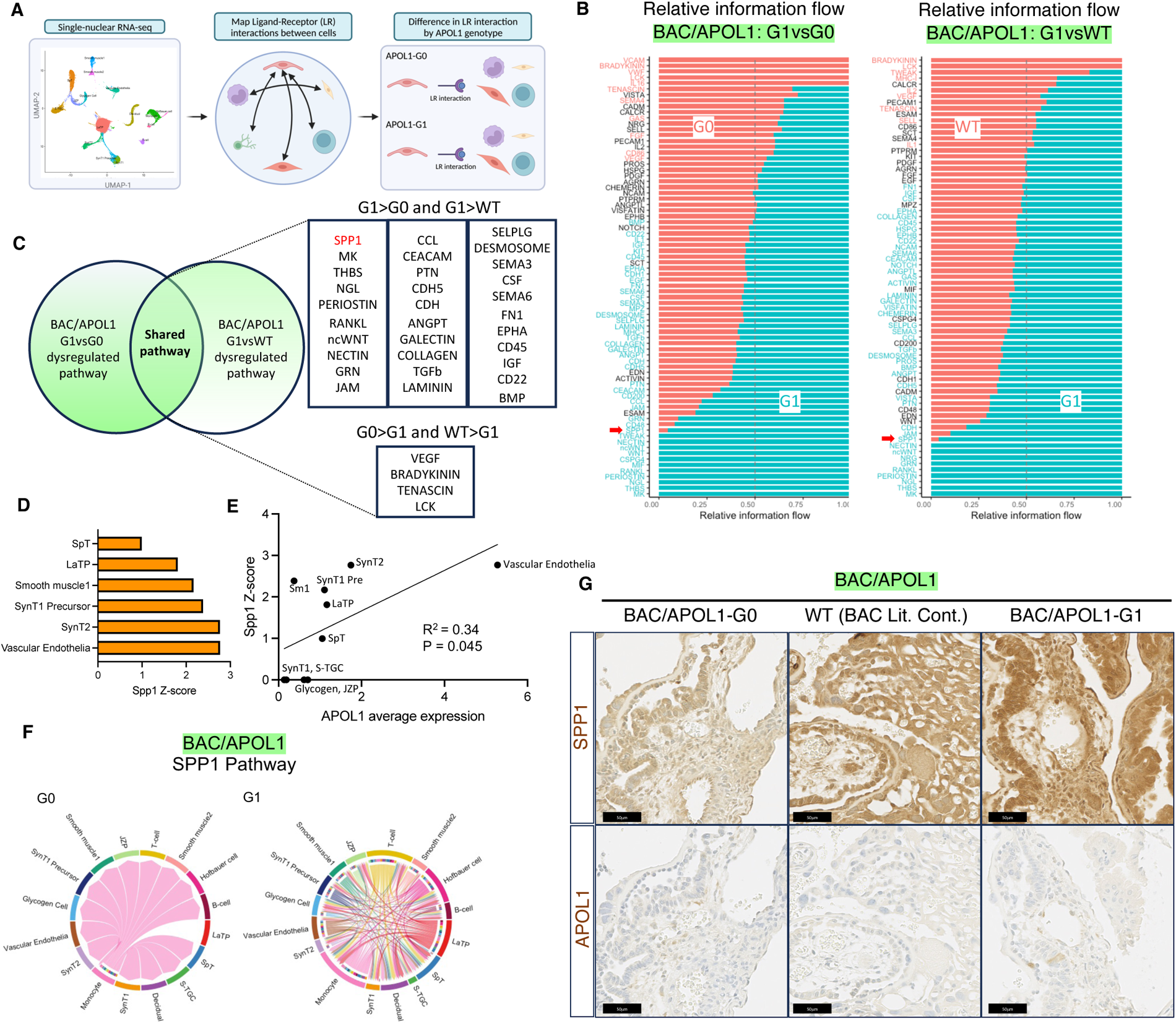
Cell-cell interaction analysis of single-nuclear RNA-seq showed inflammatory signatures in BAC/APOL1-G1 placenta including osteopontin/Spp1. (**A**) Overview of cell-cell interaction analysis using single-nuclear RNA-seq data. (**B**) Relative information flow comparing APOL1-G1 and G0 or WT (littermate control) in BAC/APOL1 model. (**C**) Venn diagram showing shared dysregulated pathways when comparing APOL1-G1 and G0 or WT (littermate control) in BAC/APOL1 model. (**D**) Activation Z-score of SPP1 pathway in each cell types comparing BAC/APOL1-G1 and BAC/APOL1-G0 placenta. (**E**) Correlational plot of each cell type in BAC/APOL1 placenta showing association between Spp1 activation Z-score and APOL1 expression, comparing APOL1-G1 and G0. (**F**) Chord plot showing more interaction of Spp1 pathway in BAC/APOL1-G1 placenta compared with BAC/APOL1-G0 placenta. (**G**) Images of immunohistochemistry showing the expression of SPP1 and APOL1 in placenta.

Cell-cell interaction analysis revealed the difference in pathways comparing APOL1-G1 placenta and APOL1-G0 or WT placenta in BAC/APOL1 model (**Figure 3B**). When we compared the dysregulated pathways in each comparison, we identified 31 shared activated and 4 shared deactivated pathways in G1 compared with G0 or WT (**Figure 3C**). With regard to ligand-receptor pairs, the osteopontin/Spp1 signaling pathway was one of the shared activated pathways, suggesting specific mechanism in BAC/APOL1-G1 (**Figure 3C**). When we checked the Spp1 activation Z-scores in each cell cluster comparing BAC/APOL1-G1 and BAC/APOL1-G0 to understand which cell type is the most likely affected by the signal, Spp1 activation Z-score was the highest in vascular endothelia, which also had the highest APOL1 expression (**Figure 3D**). Further, we observed a correlation between APOL1 expression level and Spp1 activation Z-score (**Figure 3E**). BAC/APOL1-G1 showed more Spp1 signaling between cell types than BAC/APOL1-G0 as shown in chord plot (**Figure 3F**). When we profiled up-regulated pathways from endothelial cells, Spp1 signaling pathways were prominent sending signals from endothelial cells in BAC/APOL1-G1 (**Figure S4E**). We confirmed that Spp1 detection by IHC was more pronounced in APOL1-G1 placenta (**Figure 3G, S5A**). These findings suggest that APOL1-G1 expression in vascular endothelia induced Spp1 pathway activation. Osteopontin is detected in a variety of placental cells including syncytiotrophoblasts and capillary endothelial cells, and is known to bind to integrin subunits.^23^ Osteopontin has a role in uterine vascular remodeling and decidual immune regulation.^24^

### Maternal monocytes received more signals in APOL1-G1 placenta

Based on the preeclamptic phenotype and inflammatory signal activation observed in placental single-nuclear RNA-seq in both BAC/APOL1-G1 and Alb/APOL1-G1 placenta, we hypothesized that fetal APOL1 acts on multiple cell types in placenta, including cells of maternal origin such as maternal monocytes and decidual cells. We confirmed the presence of APOL1 in plasma of dams with fetuses both with BAC/APOL1 and Alb/APOL1, albeit at low levels (**Figure S5B**). As noted in the single-nuclear RNA-seq data, monocytes and decidual cells are primarily from maternal origin, expressing *Xist*. Therefore, it is reasonable to characterize these cell types to identify the effect of APOL1 from fetuses.

When we profiled up-regulated signals to monocytes from other cell types comparing BAC/APOL1-G1 vs G0, pathways involving Cd44 as a receptor were up-regulated (**Figure 4A**). Cd44 expression in monocytes was confirmed by multiplexed FISH probing together with *Adgre1* and *Xist* (**Figure 4B**). In these maternal monocytes, the expression of *Ccl2* was upregulated in WT and G1 compared with G0 (**Figure 4B**). CCL2 is the most available chemotactic factor to decidual macrophages, and they produce a wide range of inflammatory mediators including CCL2 subsequently to create positive-feedback.^25^ Therefore, monocytes received signals from many placental cell types though Cd44 and were activated showing *Ccl2* upregulation, as indicated in chord plot showing upregulated SPP1-CD44 signaling pathway in BAC/APOL1-G1 monocytes (**Figure 4A**).

**Figure 4.**
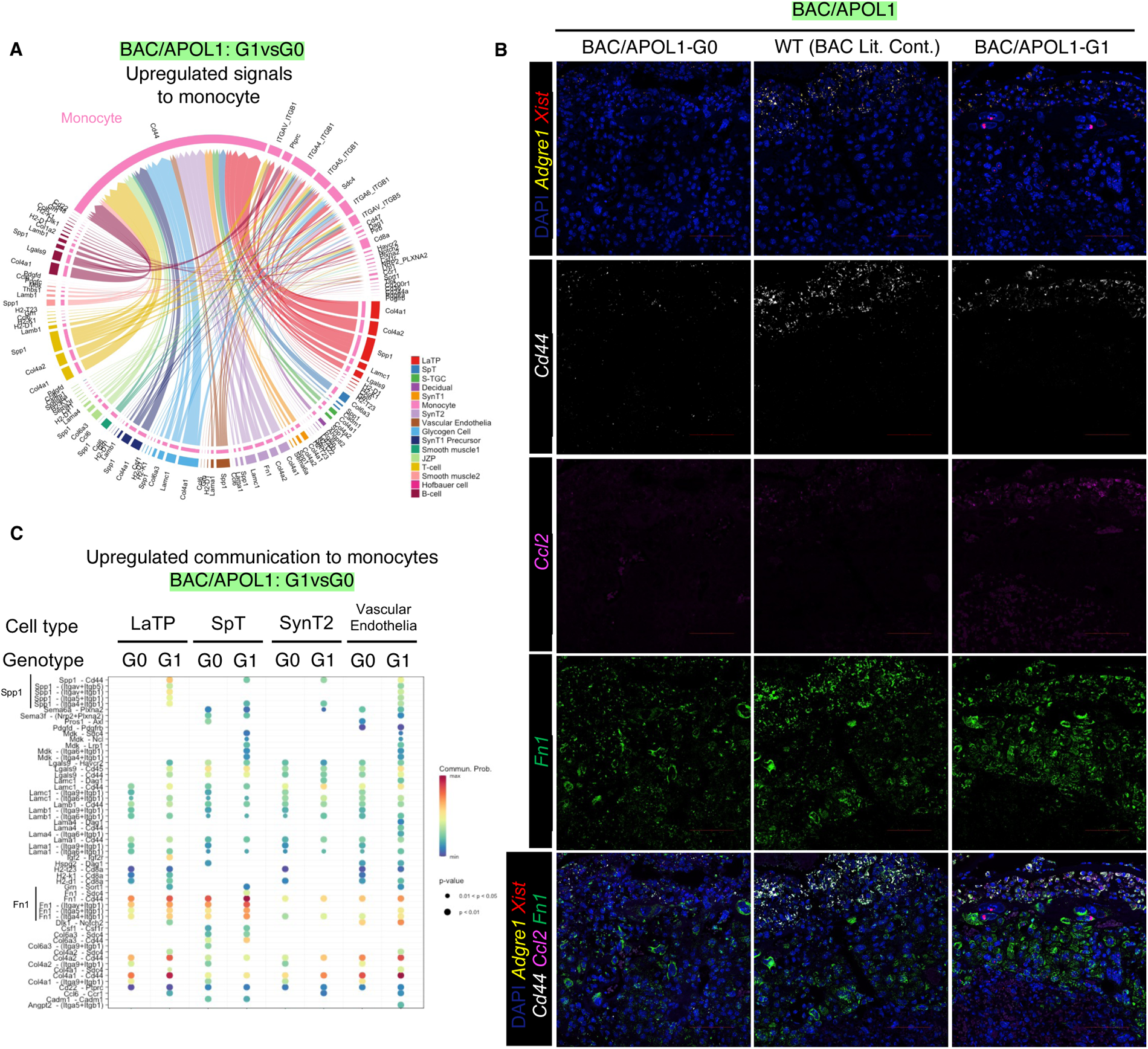
Monocytes in BAC/APOL1-G1 placenta showed more signals coming in compared with APOL1-G0 and WT placenta. (**A**) Chord plot showing upregulated signals coming to monocytes (**B**) Images of multiplex FISH showing more Cd44, Ccl2, Fn1 expression in monocytes of BAC/APOL1-G1 placenta. (**C**) Dot plot showing upregulated communications to monocytes from LaTP, SpT, SynT2 and vascular endothelial cells, comparing BAC/APOL1-G1 and G0 placenta. LaTP: labyrinth trophoblast progenitor; SpT: spongiotrophoblasts; SynT2: intermediate layer syncytiotrophoblasts

Further, we profiled upregulated signals to identify specific signals in the BAC/APOL1 model from APOL1-expressing cells to monocytes, as shown in a dot plot (**Figure 4C**). Fibronectin 1 pathway, encoded by *Fn1*, was shown to be up-regulated in labyrinth trophoblast progenitor (LaTP), SpT cells and monocytes in BAC/APOL1-G1 placenta. We observed higher expression of *Fn1* in WT and G1 compared with G0 placenta (**Figure 4B**) by FISH, concordant with single-nuclear RNA-seq data.

To evaluate the effect of circulating fetal APOL1-G1 protein on monocytes, we compared Alb/APOL1-G1 mice with WT mice (littermate controls). The results of differential interaction analysis showed that monocytes in Alb/APOL1-G1 placentas sent more signals compared with monocytes in WT placentas (**Figure 5A**). The HIF1α pathway was one of the most activated pathways (**Figure 5B**), and HIF1α was one of the activated upstream regulators (z-score 3.95) (**Figure 5C**) in monocytes of Alb/APOL1-G1 mice compared with WT mice. HIF1α activation affects signaling in monocytes and trophoblasts, upregulating cytokine RNA, including *Ccl2* and *Fn1*.^26,27^ HIF1α activation in Alb/APOL1-G1 monocytes was suggested by higher expression of *Ccl2* and *Fn1* mRNAs (**Figure 5D**). Fibronectin-1 is a glycoprotein involved in cellular adhesion and is a marker for preeclampsia.^28^ Fibronectin-1 pathway activation in both G1 and WT (littermate control) placentas in the BAC/APOL1 mice and G1 placenta in Alb/APOL1 micel indicated that activation of this pathway connected LaTP and SpT cells and monocytes, thus representing a common pathogenic mechanism in preeclamptic dams.

**Figure 5.**
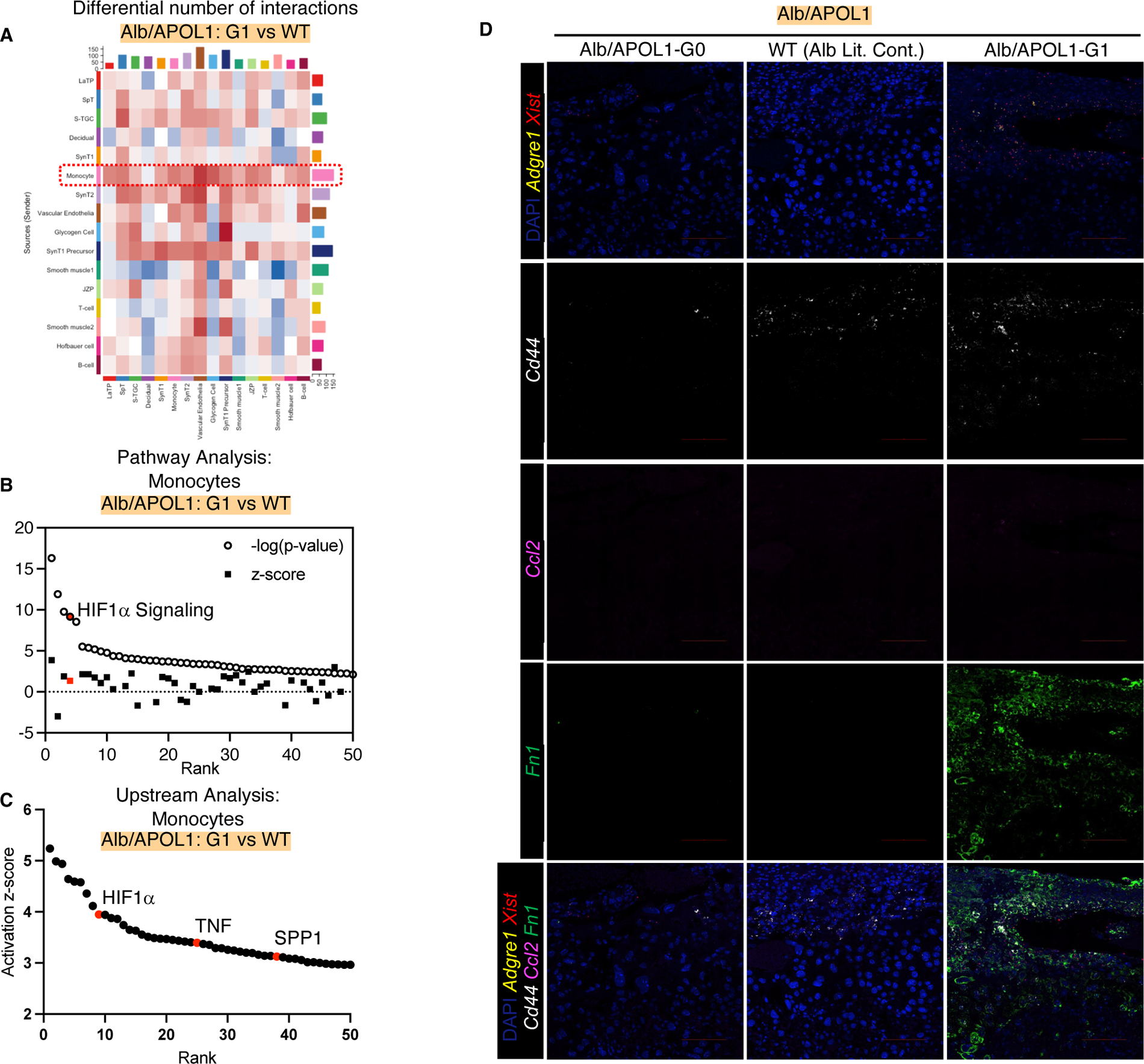
Monocytes in Alb/APOL1-G1 placenta showed activation of HIF1α signal. (**A**) Heatmap showing differential number of cell-cell interactions comparing Alb/APOL1-G1 vs WT (littermate control); red denotes more interaction in G1, blue denotes less interaction in G1 (**B**) Waterfall plot showing pathway analysis results based on monocyte DEGs comparing Alb/APOL1-G1 and WT (littermate control), showing HIF1α signaling pathway as one of the top activated pathway in Alb/APOL1-G1 (**C**) Waterfall plot showing upstream analysis results based on monocyte DEGs comparing Alb/APOL1-G1 and WT (littermate control), showing HIF1α, TNF and SPP1 as one of the top activated upstream factors in Alb/APOL1-G1 (**D**) Images of multiplex FISH showing more Cd44, Ccl2, Fn1 expression in monocytes of Alb/APOL1-G1 placenta.

## Discussion

We established and characterized two mouse models generated by *in vitro* fertilization of sperm carrying either APOL1-G0 or APOL1-G1 variants, with expression controlled by the human *APOL1* gene locus or by the mouse albumin promoter. The BAC/APOL1 IVF mouse model showed that fetal APOL1-G1 induced preeclampsia in dams, together with intrauterine fetal growth restriction of G1 fetuses. Fetal APOL1-G0 did not induce preeclampsia. Development of gestational hypertension and intrauterine growth restriction meets the diagnosis criteria of preeclampsia ^29^. These findings are congruent with previous clinical reports showing that fetal APOL1 high-risk genotype is a risk factor for preeclampsia in the mother^12–14^ and growth restriction in the infant^15^. Therefore, this BAC/APOL1-G1 IVF mouse model was a relevant mouse model to study the molecular mechanism of preeclampsia, especially preeclampsia associated with fetal APOL1-G1 induced one. It is also worth noting that Alb/APOL1-G1 model showed similar level of phenotypes compared with BAC/APOL1-G1 model, except for change in the of sFlt-1/PlGF-2 ratio. The limited APOL1 expression in liver and plasma in Alb/APOL1 model^18^ and genetic background could be the reasons for the difference with BAC/APOL1 model.

Preeclampsia mouse models reported so far can be categorized as either a mixed strain approach^30,31^, germline transgenic approach^32^, exogenous agent approach^33^, surgical approach^34^, and viral approach.^2,27,35^ The mice reported here are unique in that the transgene is only in fetuses, due to the use of IVF.

The only previous APOL1 transgenic mouse that has shown preeclampsia is described by Bruggeman *et al.* and is *Nphs1* promotor driven APOL1-G0 and APOL1-G2 transgenic mice; these mice develop preeclampsia in both -G0 and -G2 genotypes^16^. The Bruggeman *et al.* study mainly characterize dams carrying the transgene, but also implicate fetal effect by the observation that WT FVB/N females used as dams for founder males exhibit seizures or sudden death during pregnancy. Wu *et al.* showed that the APOL1-G2 variant, when over-expressed in endothelial cells. induced endothelial inflammation and dysfunction, leading to sepsis.^36^ As the BAC/APOL1-G1 placenta in the present study showed the highest expression of APOL1 in endothelial cells, it is reasonable to speculate that endogenous APOL1-G1 induced endothelial inflammation and dysfunction.

Due to the complexity and dynamics of fetal-maternal interplay involving multiple cell types during pregnancy, it has been a challenge to characterize cell-specific signals. Placental single-nuclear RNA-seq showed dysregulated genes and pathways at cellular resolution comparing 1) APOL1-G1 vs APOL1-G0 and 2) APOL-G1 vs wild-type littermate control.

The osteopontin/Spp1 pathway was activated in multiple cell types in APOL1-G1 placenta, most highly in vascular endothelial cells expressing high APOL1. A previous report showed that plasma osteopontin concentrations are increased in preeclamptic patients with extensive endothelial injury.^37^ Osteopontin activates trophoblast invasion in *in vitro*.^23^ We characterized mice at day E18.5, which is not when the active stage of vascular invasion occurs. Nevertheless, as indicated in a human study^37^, high levels of plasma osteopontin may be a marker of irregular vascular invasion and preeclampsia.

Cell-cell interaction analysis indicated more communication in BAC/APOL1-G1 placenta, mediated by maternal monocytes showing expression of Cd44. The Spp1 signaling pathway was activated in both BAC/APOL1-G1 and Alb/APOL1-G1placentas, compared with G0 placenta. This indicates that circulating APOL1-G1 may induce activation of monocytes and/or other placental cells. It is interesting to note that Alb/APOL1-G1 vascular endothelial cells showed activation of the SPP1 pathway compared with Alb/APOL1-G0, albeit at a lower level. Fetal-maternal mismatch of *APOL1* genotype has been reported to be a risk factor for preeclampsia.^14^

The role of maternal monocytes and macrophages in preeclampsia has been investigated in human placenta tissue and in rat models.^38,39^ Maternal monocytes become activated during their passage through the placenta.^40^ APOL1 was detected in plasma of dams with both BAC/APOL1 and Alb/APOL1 fetuses, although at a low level. The limited APOL1 expression in plasma in the Alb/APOL1 model enabled us to investigate the effect of circulating APOL1 on maternal monocytes. HIF1α is a well-characterized pathogenetic molecule in preeclampsia.^41^ In HIF1α transgenic mouse models, both systemically-expressed ^42^ and trophoblast-expressed ^27^ HIF1α showed preeclampsia. Here we identified for the first time HIF1α pathway activation in monocytes in Alb/APOL1-G1 placenta. Further, we observed common pro-inflammatory gene upregulation, including MCP1/*Ccl2* in both BAC/APOL1-G1 and Alb/APOL1-G1 placentas. These findings suggest that fetal-derived circulating APOL1-G1 induced preeclampsia in the dams, associated with maternal monocytes activation.

This study focused only on differences between APOL1-G0 and APOL1-G1 variants with regard to preeclampsia; we did not study APOL1-G2, as this line has lower APOL1-G2 expression. We did not observe obvious histological differences in placenta. One of the possible reasons for this result is that mice do not have deep trophoblast invasion into the uterine wall as humans or rats do.^43,44^ We also did not observe other preeclampsia phenotypes, such as elevated serum creatinine, albuminuria and elevated liver enzymes (**Supplemental Figure 1**). These models have only a fetal factor promoting preeclampsia, *e.g. APOL1-G1* genotype, and no maternal factors. In patients, other factors may contribute to severe preeclampsia, in addition to fetal *APOL1* high-risk genotype.

The BAC/APOL1 mouse background is 129Svj, so the genetic background of fetuses in BAC/APOL1 IVF model are all FVB/N/129Svj. On the other hand, the Alb/APOL1 mice background is FVB/N, so the genetic background of fetuses in Alb/APOL1 IVF model are all FVB/N. Because the genetic backgrounds may affect the biology of the two mouse models, direct comparisons were made within each model, comparing APOL1-G1 and APOL1-G0 or APOL1-G1 and wild-type (APOL1-G1 littermate control).

In conclusion, we characterized two mouse models involving IVF using sperm carrying the APOL1-G1 variant; in both models, dams manifested preeclamptic phenotypes. Further investigation of this disease process is warranted to identify novel therapeutic targets for APOL1-associated preeclampsia.

## Methods

### Sex as a biological variable

Our study exclusively examined female mice for dams because the disease modeled is only relevant in females. Further, our study examined male and female animals for fetuses, and similar findings are reported for both sexes.

### Mice

Mice in experimental groups were matched for sex and age. Mice were housed in cages on a constant 12-h light/dark cycle, with controlled temperature and humidity and food and water provided ad libitum. Sample sizes for experiments were determined without formal power calculations.

We used human APOL1 gene locus transgenic mice (BAC/APOL1 mice)^17,45^ with APOL1-G0 and APOL1-G1 heterozygotes (Merck & Co., Inc. (Rahway, NJ, USA)), and mice carrying the human APOL1 gene under the control of the mouse albumin promoter (Alb/APOL1 mice)^18^ with APOL1-G0 and APOL1-G1 heterozygotes to obtain sperm for *in vitro* fertilization (IVF). Sperm from both transgenic lines was used for *in vitro* fertilization to oocytes from FVB/N mice, then 2-cell stage embryos were transferred to CD-1 female recipients. From the CD-1 dams, 24-h urine collections were obtained from gestational days 17 to 18, using metabolic cages.

Tissues were obtained at gestational day 18.5 (E18.5). Mice were anesthetized mice with 2, 2, 2-trimbromoethanol (Avertin, 2,2,2-tribromoethanol, Sigma-Aldrich, St. Louis, MO). Maternal plasma, kidneys, placentas and fetuses were collected.

Tails of fetuses were submitted to Transnetyx (Cordova, TN) for genotyping for APOL1-G0 or G1 and sex determination. Fetuses and placentas were measured and weighed. Half the placental samples and fetuses were fixed with 10% buffered formalin for 24 h and paraffin embedded. One fourth of each placenta were snap frozen by liquid nitrogen and kept in -80C° for single-nuclear RNA-seq.

### Blood pressure measurement

Tail blood pressure was measured in the dams at gestational day 18 using CODA non-invasive blood pressure system (Kent Scientific, Torrington, CT). We accustomed mice to the device for one to two days before measurement. After acclimation for at least 10 min on the measurement setup on a 37C° warming tray, systolic blood pressure was measured by 25 cycles with 5 seconds between cycles. Median values were calculated.

### Mouse chemistry measurements

Urinary albumin levels were measured with the Albuwell M (mouse albumin ELISA) (1011, Ethos Biosciences, Logan Township, NJ). We measured the urine creatinine concentration with Creatinine Companion kit (1012, Ethos Biosciences). Albuminuria was reported as the ratio of urinary albumin to creatinine. Plasma creatinine was measured by isotope dilution LC-MS/MS at The University of Alabama at Birmingham O’Brien Center Core C (Birmingham, AL). Plasma sFlt1 and PlGF-2 levels were measured by Mouse VEGFR1/Flt-1 Quantikine ELISA Kit and Mouse PlGF-2 Quantkine ELISA Kit (MVR100 and MP200, Biotechne, Minneapolis, MN). All procedures were performed in accordance with the manufacturers’ protocols.

### Single-nucleus RNA-sequencing

Nuclei from each one placenta of male offspring (BAC/APOL1: G0, G1, WT (G1 littermate control); Alb/APOL1: G0, G1, WT (G1 littermate control)) were prepared as follows. Snap frozen placenta were lysed in EZlysis buffer (#NUC101-1KT, Sigma, Darmstadt, Germany) after dissecting decidua and homogenized 30 times using a loose Dounce homogenizer and 5 times in a tight pestle. After 5 min incubation, the homogenate was passed through a 40 µm filter (43-50040, PluriSelect, El Cajon, CA) and centrifuged at 500g at 4°C for 5 min. The pellet was washed with EZlysis buffer and again centrifuged at 500g at 4°C for 5 min. The pellet was resuspended with DPBS with 1% FBS to make final nuclei prep for loading on to a 10x Chromium Chip G (10X Genetics, Pleasanton, CA) for formation of gel beads in emulsion (GEM).

Single nuclear isolation, RNA capture, cDNA preparation, and library preparation were performed following the manufacturer’s protocol (Chromium Next GEM Single Cell 3’ Reagent Kit, v3.1 chemistry, 10x Genomics). Prepared cDNA libraries were sequenced at DNA Sequencing and Genomics Core (NHLBI, NIH). The sequencing and mapping statistics of single-nucleus RNA-seq are in **Supplemental Table S1**.

### Single-nuclear RNA-seq analysis

Preprocessing was performed with the Cell Ranger v 6.1.2 software (10x Genomics) using the default parameters. The reference was built from mm10 reference genome, supplemented with human APOL1 sequences. Removal of ambient RNA was conducted by SoupX (version 1.5.2)^46^ following the default protocol by “autoEstCont” and “adjustCounts” functions. After removal of ambient RNA, integration of single-nucleus gene expression data was performed using Seurat (version 4.0.5) and SeuratData (version 0.2.2)^47^ after filtering out nuclei with any of the following features: total RNA count < 500 or > 4000 or percentage of mitochondrial transcripts > 25%. After filtering, 38351 nuclei remained for downstream analysis. Clustering was performed by constructing a K-nearest neighbor graph by first 20 principal components and applying the Louvain algorithm with resolution 0.5. Dimensional reduction was performed with UMAP, 24 clusters were annotated based on expression of specific markers.^19^ Differential expressed genes among cell types were assessed with the Seurat FindMarkers function with default parameters. DEG were identified for each paired comparison of genotype, using a cut-off of adjusted P <0.05. Pathway analysis was performed using QIAGEN Ingenuity Pathway Analysis (IPA) software^48^, using DEG sets as inputs. Analysis match function of IPA was used to perform comparative analysis with data from GSE143953^22^ and GSE93839^21^. Cell-cell interaction analysis was performed using CellChat (version 1.5.0)^49^ with mouse protein-protein interactions loaded.

### Pathological evaluation

FFPE mouse placenta and kidney tissue sections, cut at 4 µm, were stained with hematoxylin and eosin, periodic acid-Schiff (PAS) reagents for routine histological assessment.

### Fluorescent in situ hybridization (FISH)

Fluorescent *in situ* detection of mRNA was performed on tissue sections from the mouse formalin-fixed paraffin-embedded (FFPE) blocks using the RNAscope Hiplex12 Reagents Kit (488, 550, 650) v2 (#324419, Advanced Cell Diagnostics, Biotechne, Minneapolis, MN). Briefly, 5 μm FFPE tissue sections were de-paraffinized, boiled with RNAscope Target Retrieval Reagent for 15 min, and then protease digested at 40°C for 30 min, followed by hybridization for 2 h at 40°C with probes. RNA probe-Hs-APOL1-No-XMm (#459791), Mm-Ccl2 (#311791), Mm-Fn1 (#316951), Mm-Adgre1 (#460651), Mm-Cd44 (#476201), Mm-Xist (#418281) were used for hybridization. Specific probe binding sites were visualized following RNAscope Hiplex12 Reagents Kit (488, 550, 650) v2 (Advanced Cell Diagnostics, Biotechne) (#324419).

### Immunohistochemistry

FFPE tissue sections were deparaffinized and rehydrated. Antigen retrieval was performed by heating in citrate-buffered medium for 15 min in a 99°C hot water bath. Tissues were blocked by 2.5% normal horse serum. Sections were incubated for one h at room temperature with 5 μg/ml of primary antibody for APOL1 (5.17D12, rabbit monoclonal) that was kindly provided by Genentech (South San Francisco, CA)^50^ or for osteopontin (SPP1) (LF-175, Kerafast, Boston, MA). Sections were processed following ImmPRESS HRP Universal Antibody (horse anti-mouse/rabbit IgG) Polymer Detection Kit and ImmPACT DAB EqV Peroxidase (HRP) Substrate (Vector Laboratories, Burlingame, CA) protocol, and counter-stained with hematoxylin.

### Confocal microscopy

A Yokogawa CSU-W1 SoRa spinning disk confocal scanhead (Yokogawa, Sugar Land, TX), with 50-micron pinhole (standard confocal mode, no SoRa), mounted on a Nikon Ti2 microscope running NIS-elements 5.21.02 software (Nikon Instruments, Melville, NY), was used to collect tiles of multi-color fluorescence images. Fluorescence image channels were obtained sequentially, while sharing the Yokogawa T405/488/568/647 dichroic. For DAPI fluorescence: excited by the 405nm laser, emission was filtered by ET455/58 (Chroma, Technology Corp, Bellows Falls, VT). For green fluorescence: excited by the 488nm laser, emission filtered by ET520/40 (Chroma). For orange fluorescence: excited by the 561nm laser; emission filtered by ET605/52 (Chroma). For far red fluorescence: excited by the 640nm laser; emission filtered by ET655LP (Chroma). Images used for quantitation of stain prevalence were acquired with the Nikon Apo TIRF 60x/1.49 Oil DIC N2 objective lens, producing a confocal section thickness of 1.2 microns for all fluorescence channels.

### Statistics

We conducted t-test for two-group comparisons and one-way ANOVA for comparisons involving more than two groups, with multiple test correction. A P value less than 0.05 was considered significant.

### Study approval

All mouse experiments were conducted in accordance with the National Institutes of Health Guide for the Care and Use of Laboratory Animals and were approved in advance by the NIDDK Animal Care and Use Committee (Animal study proposal, K097-KDB-17 & K096-KDB-20).

### Data availability

Original data files and count tables have been deposited in GEO. Other data are available from the authors upon request.

## Author Contributions

TY, AZR and JBK conceived the study design. TY, SS and HL conducted mouse experiments. TY conducted single-nuclear RNA-seq capture, TY and KZL conducted single-nuclear RNA-seq data analysis. KZL and YZ supported sequencing analysis. AZR and SR assessed pathology. TY drafted the manuscript and all the authors contributed for edits.

## Acknowledgements

We thank the DNA Sequencing and Genomics Core (NHLBI/NIH) for sequencing support, Dr. Luis Menezes (NIDDK/NIH) for critical manuscript review. We appreciate receiving BAC/APOL1 mice from Merck & Co., Inc., Rahway, NJ, USA. We appreciate receiving APOL1 antibodies from Genentech, South San Francisco, CA. Figures were created with Morpheus (https://software.broadinstitute.org/morpheus) and BioRender.com. This work utilized the resources of the NIH HPC Biowulf cluster (http://hpc.nih.gov) and NIDDK Advanced Light Microscopy & Image Analysis Core (ALMIAC) and NIDDK Mouse Transgenic Core Facility. The content of this publication does not necessarily reflect the views or policies of the Department of Health and Human Services, nor does mention of trade names, commercial products, or organizations imply endorsement by the U.S. Government.

## Funding

This project has been funded in part with federal funds from the National Cancer Institute, National Institutes of Health, under contract 75N91019D00024. This work was supported by the Intramural Research Program of the NIH, including the National Cancer Institute, Center for Cancer Research and the NIDDK (Project 1ZIADK043308-28). KJR received support from the Preeclampsia Foundation.

---

**Supplemental Figure 1.**
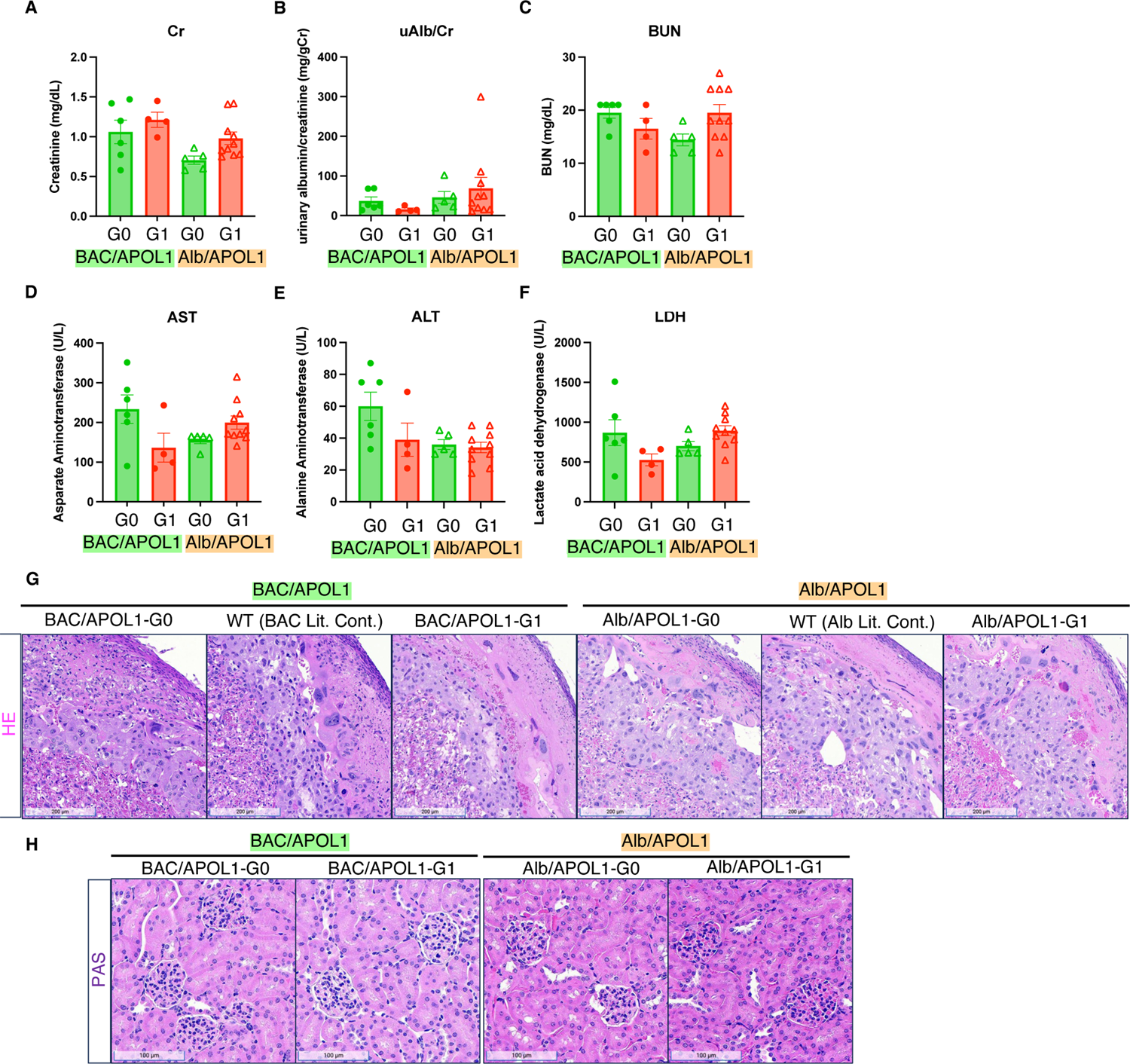
Additional phenotypes and pathologies of BAC/APOL1 and Alb/APOL1 models. (**A-F**) Serum creatinine, urinary albumin-to-creatinine ratio, blood urea nitrogen, AST, ALT, and LDH of dams identified by fetal genotype (**G**) Representative images of HE staining of placentas (**H**) Representative images of PAS staining of maternal kidneys

**Supplemental Figure 2.**
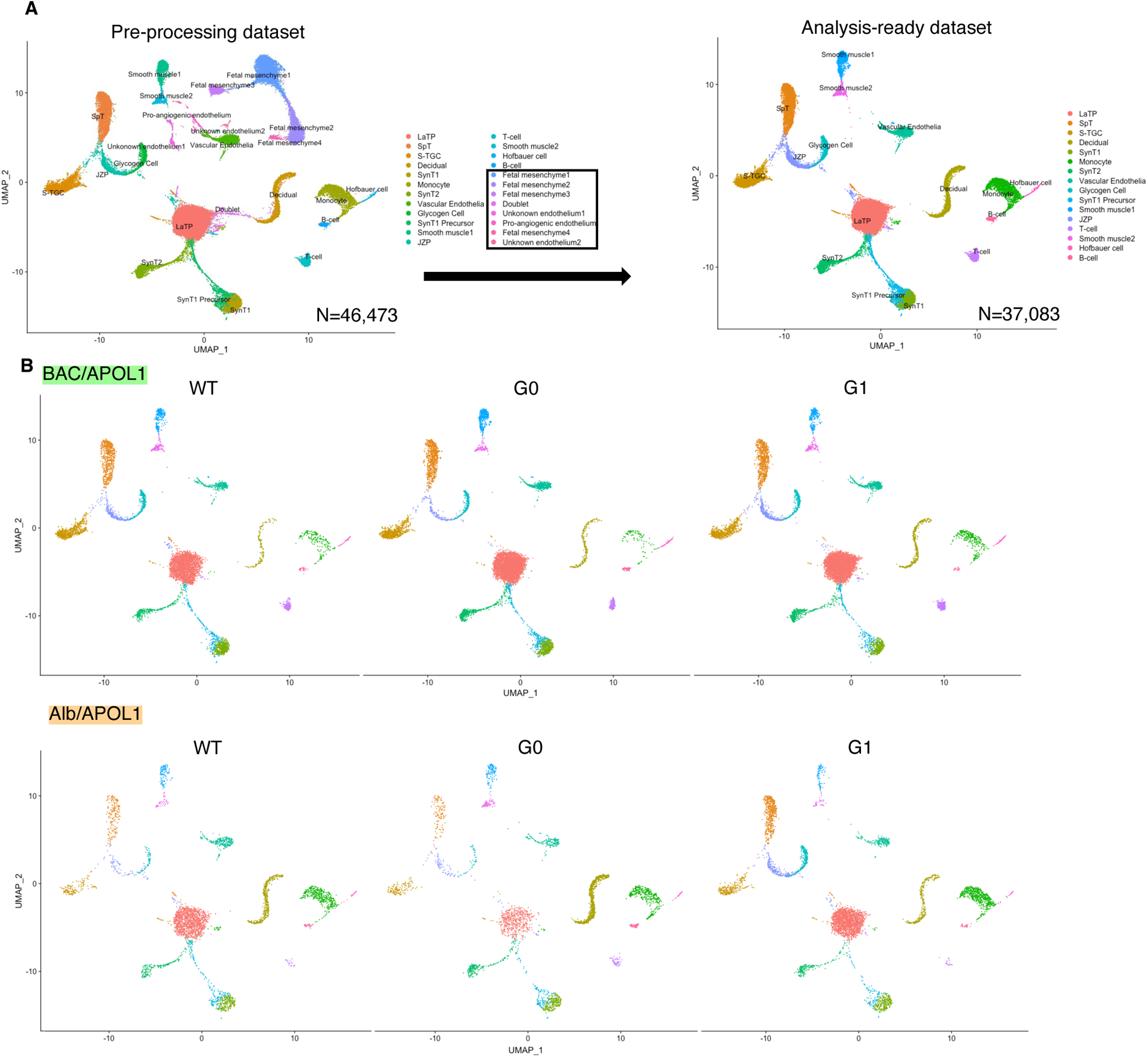
Supplemental single-nuclear RNA-seq data, pre-processing and UMAP. (**A**) UMAP plot of pre-processing dataset (N=46,473) and analysis-ready dataset (N=37,083) (**B**) UMAP plots separated by each model

**Supplemental Figure 3.**
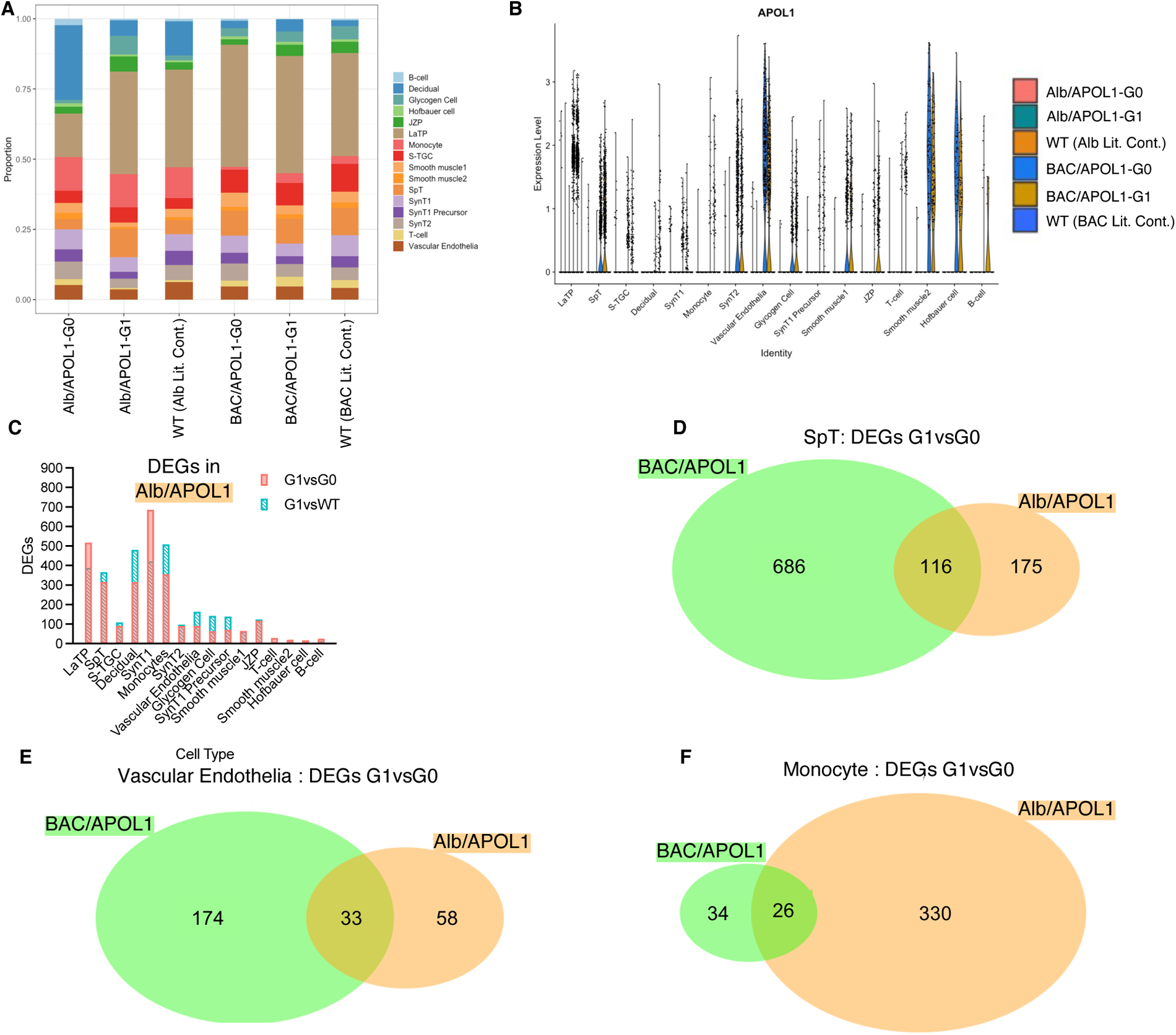
Additional single-nuclear RNA-seq data. (**A**) Ratio of nuclei grouped to each cluster by each sample (**B**) Violin plot showing APOL1 expression in each sample in each cluster (**C**) Number of DEGs in each cluster, comparing APOL1 genotype (G1 vs G0 and G1 vs WT) in Alb/APOL1 model. (**D-F**) Venn diagram of DEGs of SpT, vascular endothelia, and monocyte, comparing APOL1 genotype (G1 vs G0) in both BAC/APOL1 and Alb/APOL1 model

**Supplemental Figure 4.**
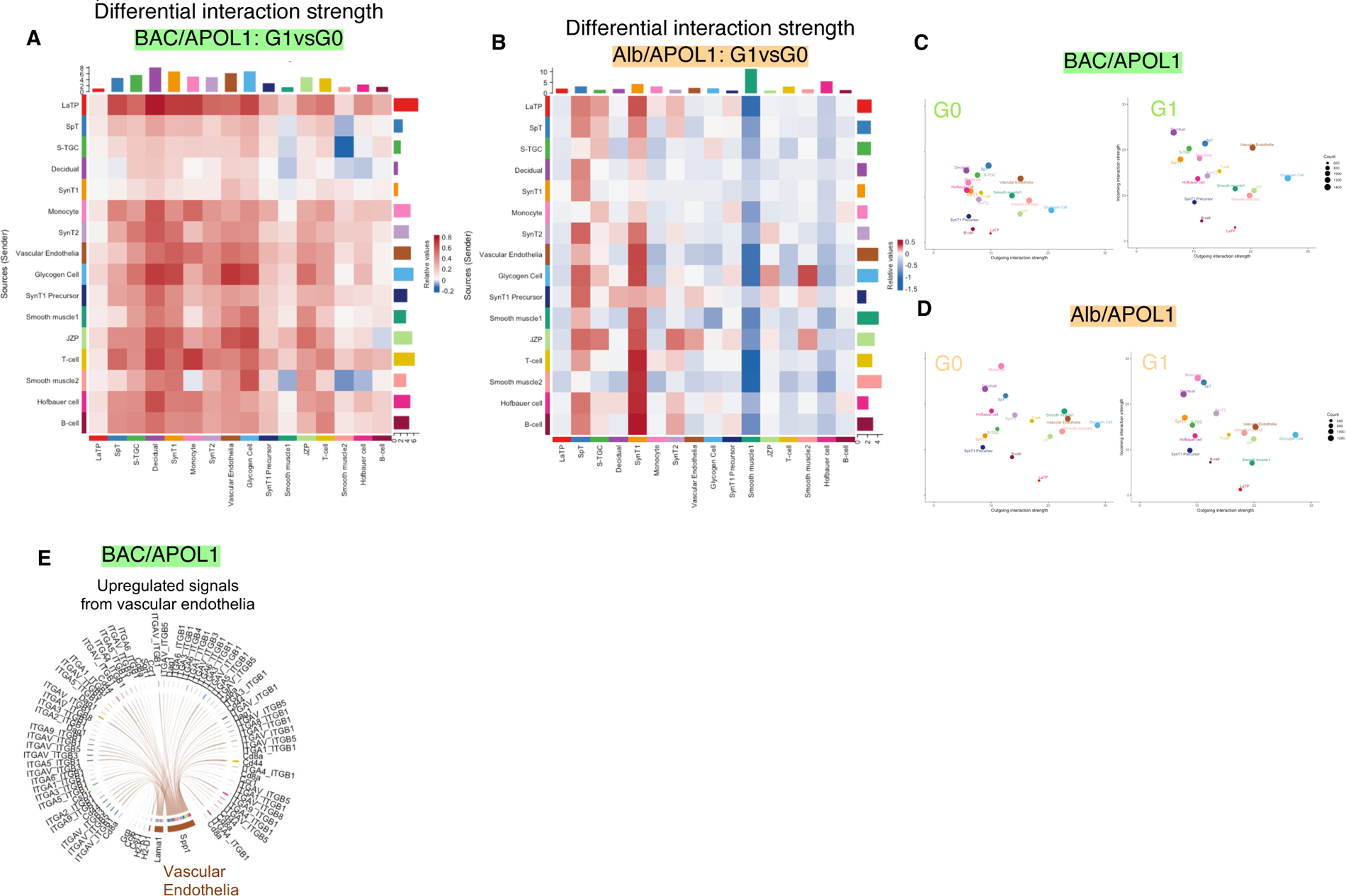
Additional cell-cell interaction analysis data. (**A,B**) Heatmaps showing differential interaction strength among cell types comparing APOL1-G1 vs G0 in each of BAC/APOL1, Alb/APOL1 models (**C,D**) 2D plot showing incoming and outgoing interaction strength in each cell type comparing APOL1-G1 vs G0 in each of BAC/APOL1, Alb/APOL1 models (**E**) Chord plot showing upregulated signals from vascular endothelia in BAC/APOL1 model

**Supplemental Figure 5.**
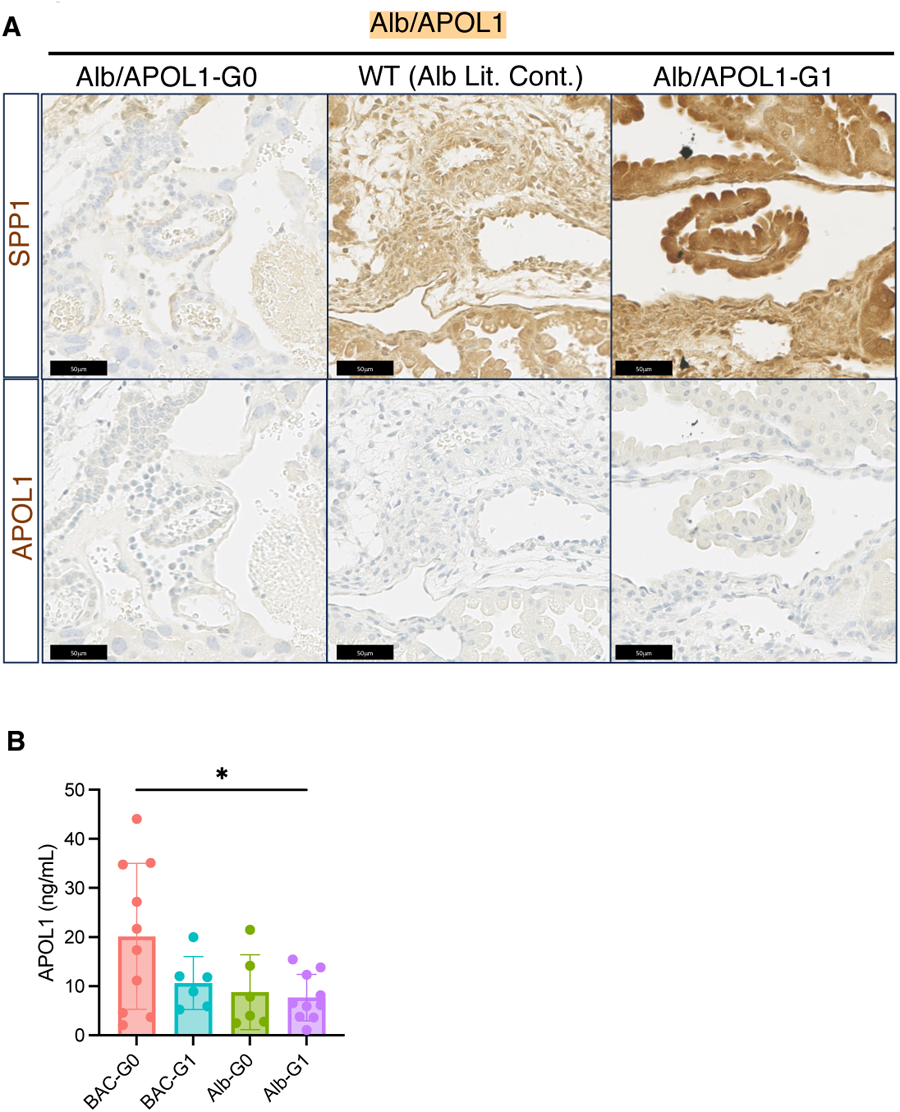
Additional IHC of placenta from Alb/APOL1 model and plasma APOL1 level of dams. (**A**) Representative IHC images of SPP1 and APOL1 of Alb/APOL1 model placenta (**B**) Plasma APOL1 level measured in dams

## Notes

### Competing Interest Statement

The authors have declared no competing interest.

### Summary of Updates

The title was updated to "Preeclampsia in mice carrying fetuses with APOL1 risk variants ".

