## Supplemental Figures for "Preeclampsia in mice carrying fetuses with APOL1 risk variants"

Figure S1

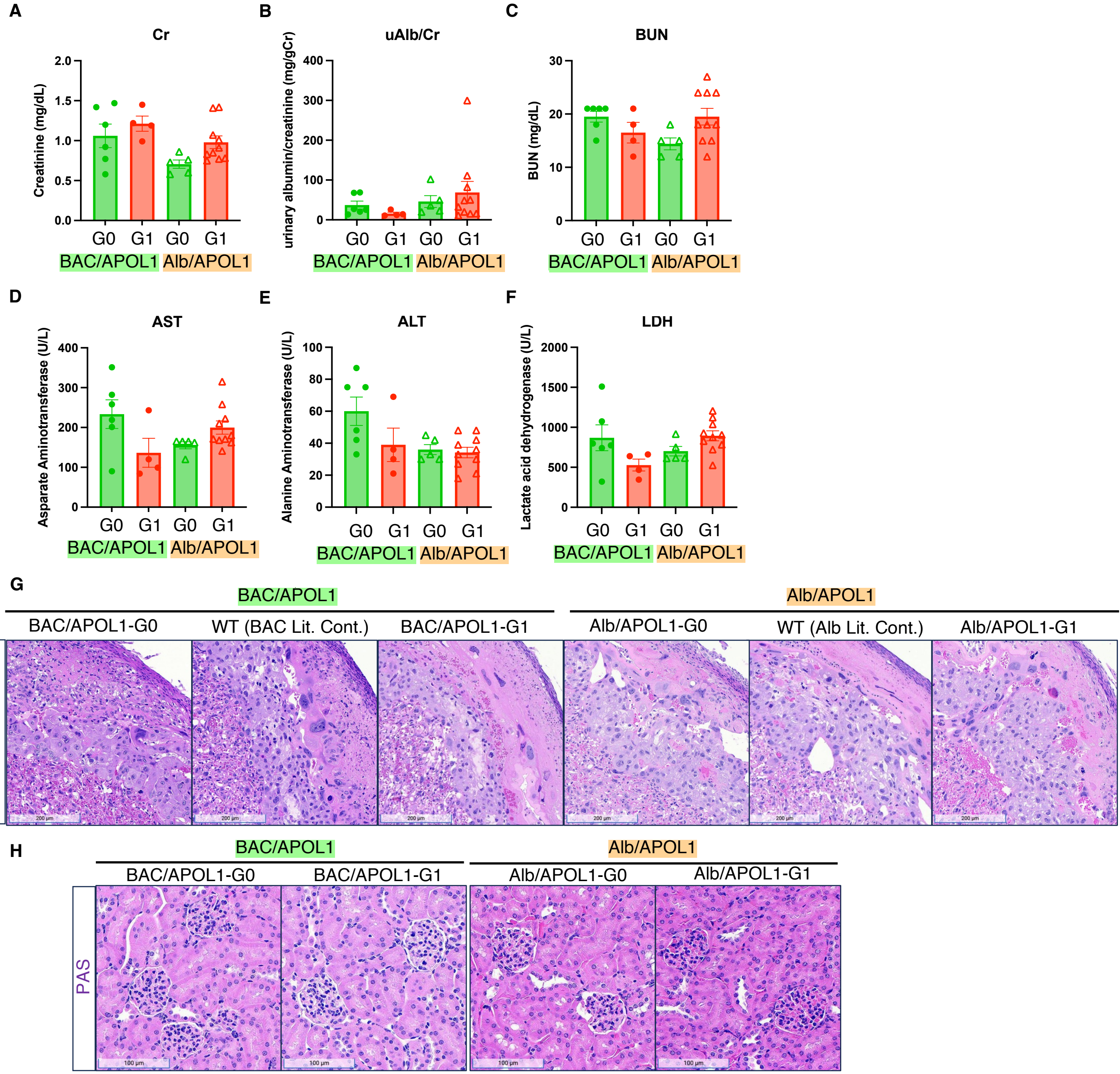


**Supplemental Figure 1. Additional phenotypes and pathologies of BAC/APOL1 and Alb/APOL1 models**
(**A-F**) Serum creatinine, urinary albumin-to-creatinine ratio, blood urea nitrogen, AST, ALT, and LDH of dams identified by fetal genotype
(**G**) Representative images of HE staining of placentas (**H**) Representative images of PAS staining of maternal kidneys

**Figure S2**

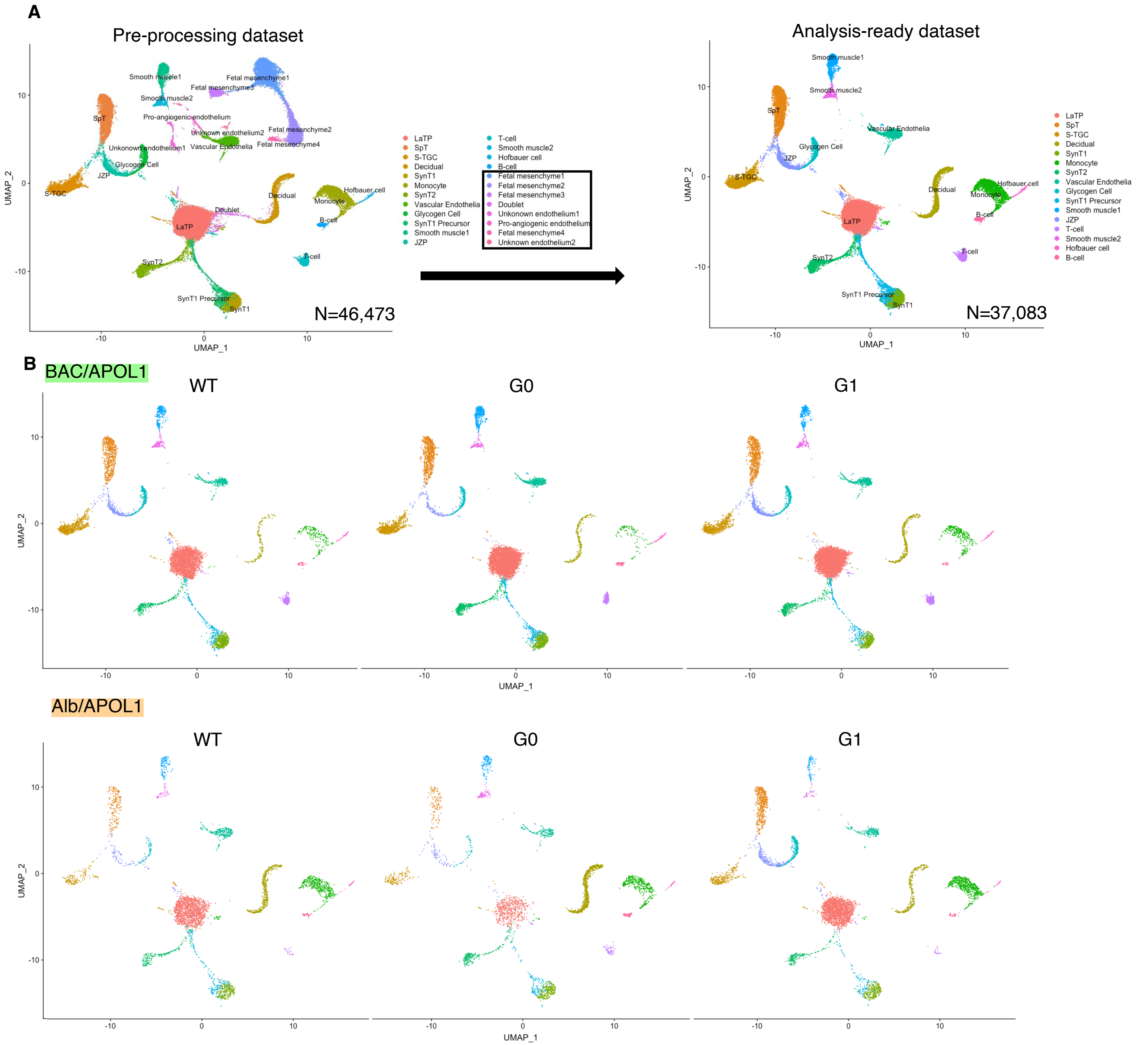


**Supplemental Figure 2. Supplemental single-nuclear RNA-seq data, pre-processing and UMAP**
(**A**) UMAP plot of pre-processing dataset (N=46,473) and analysis-ready dataset (N=37,083) (**B**) UMAP plots separated by each model

**Figure S3**

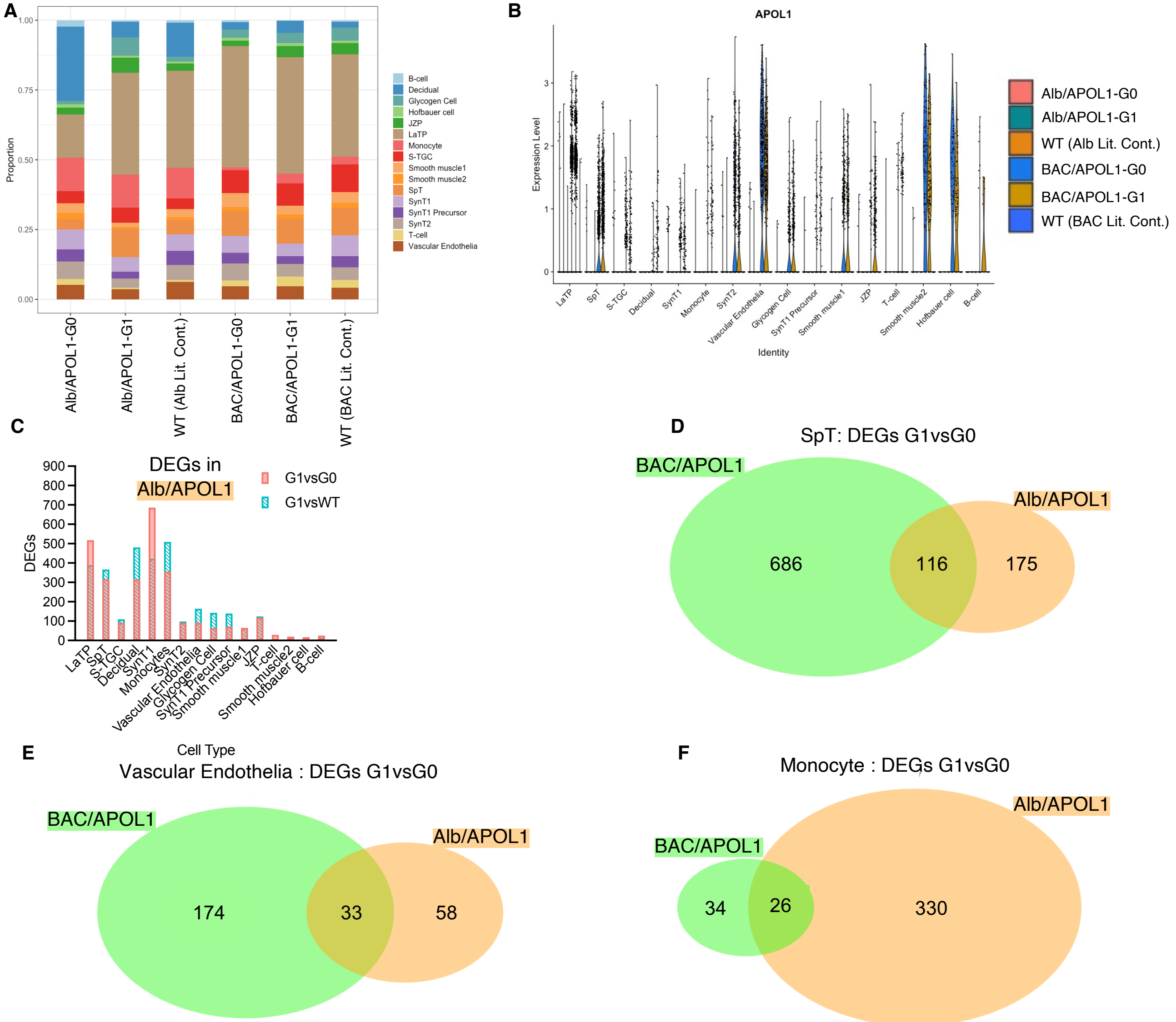


**Supplemental Figure 3. Additional single-nuclear RNA-seq data**
(**A**) Ratio of nuclei grouped to each cluster by each sample (**B**) Violin plot showing APOL1 expression in each sample in each cluster
(**C**) Number of DEGs in each cluster, comparing APOL1 genotype (G1 vs G0 and G1 vs WT) in Alb/APOL1 model.
(**D-F**) Venn diagram of DEGs of SpT, vascular endothelia, and monocyte, comparing APOL1 genotype (G1 vs G0) in both BAC/APOL1 and Alb/APOL1 model

**Figure S4**

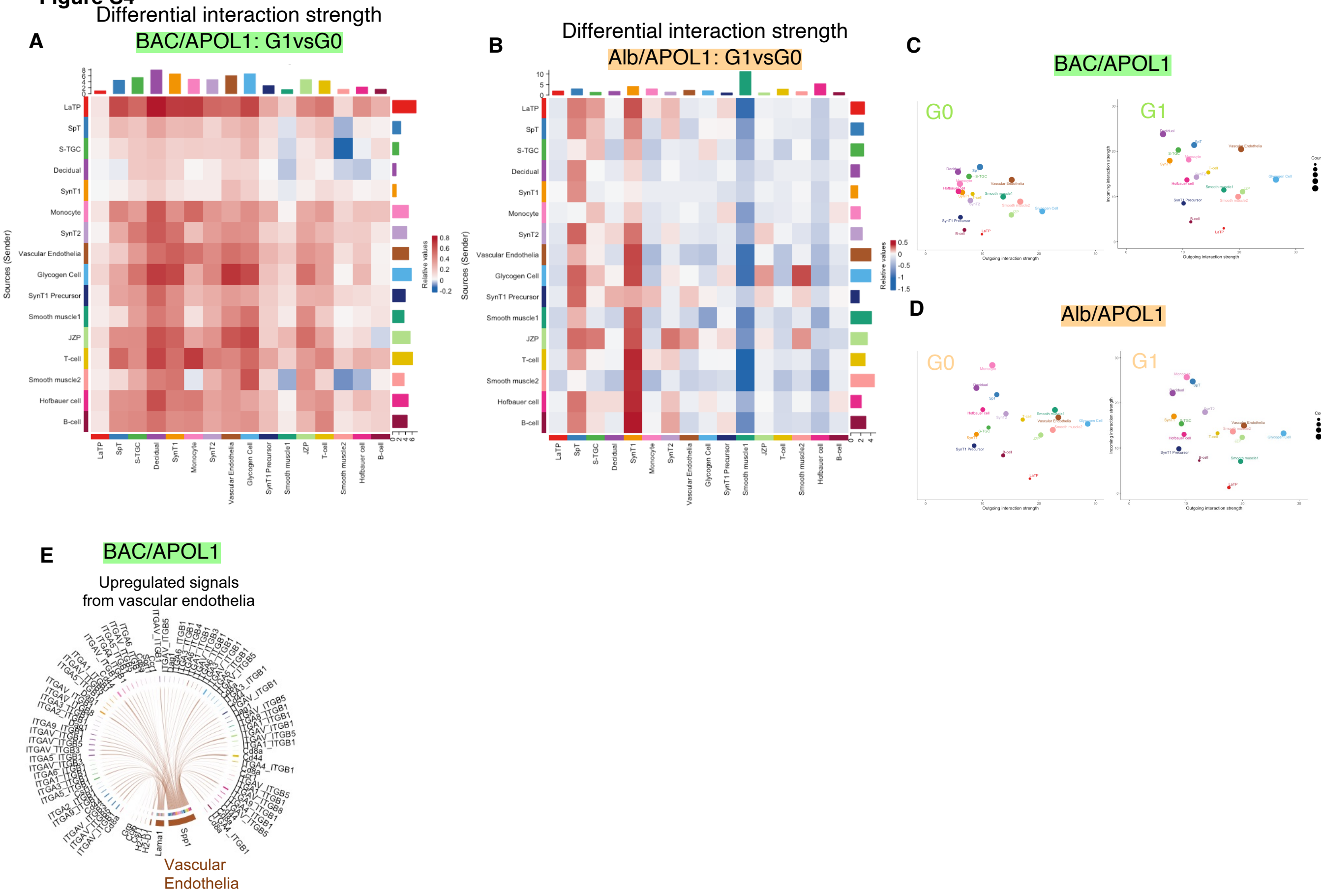


**Supplemental Figure 4. Additional cell-cell interaction analysis data**

(**A,B**) Heatmaps showing differential interaction strength among cell types comparing APOL1-G1 vs G0 in each of BAC/APOL1, Alb/APOL1 models

(**C,D**) 2D plot showing incoming and outgoing interaction strength in each cell type comparing APOL1-G1 vs G0 in each of BAC/APOL1, Alb/APOL1 models

(**E**) Chord plot showing upregulated signals from vascular endothelia in BAC/APOL1 model

**Figure S5**

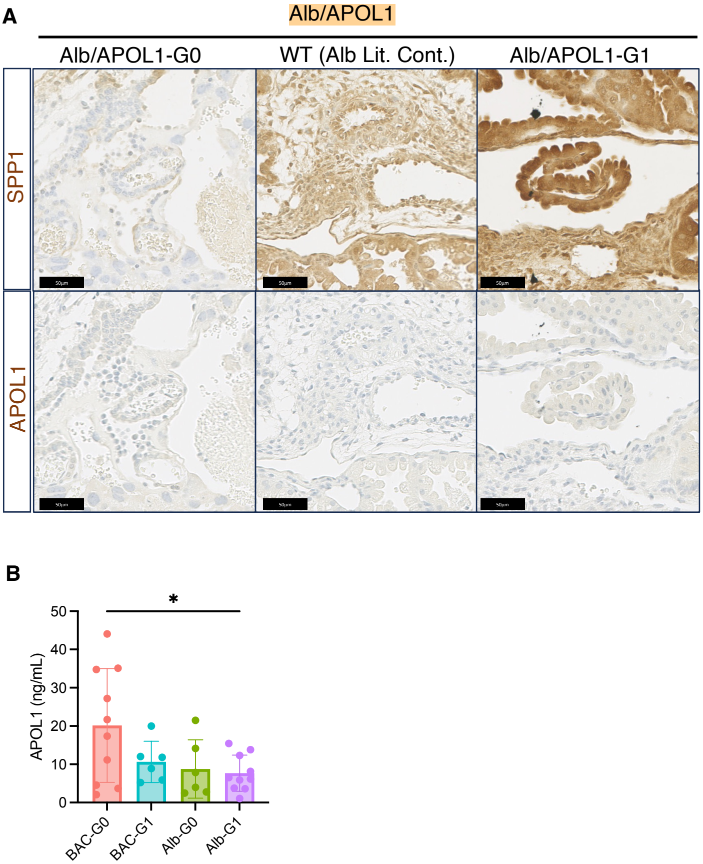


**Supplemental Figure 5. Additional IHC of placenta from Alb/APOL1 model and plasma APOL1 level of dams.**
(**A**) Representative IHC images of SPP1 and APOL1 of Alb/APOL1 model placenta (**B**) Plasma APOL1 level measured in dams
